# Higher concentrations of bacterial enveloped virus Phi6 can protect the virus from environmental decay

**DOI:** 10.1101/2021.05.17.444592

**Authors:** Ronald Bangiyev, Maxim Chudaev, Donald W. Schaffner, Emanuel Goldman

## Abstract

Phage Phi6 is an enveloped virus considered as a possible non-pathogenic surrogate for SARS-CoV-2 and other viral pathogens in transmission studies. Higher input amounts of bacteriophage Phi6 are shown to delay and protect the phage from environmental decay, both when the phage are dried in plastic tubes, and when they are stored in saline solution at 4°C. When bacteriophage Phi6 are placed in LB (Luria-Bertani) growth medium prior to placement on the plastic surface, viral recovery is not influenced by the starting concentration. The protection is reflected in longer half-lives of the phage at higher concentrations compared to lower. Because experiments supporting the possibility of fomite transmission of SARS-CoV-2 and other viruses rely upon survival of infectious virus following inoculation of various surfaces, high initial amounts of input virus on a surface may generate artificially inflated survival times compared to realistic lower levels of virus that a subject would normally encounter. This is not only because there are extra half-lives to go through at the higher concentrations, but also because the half-lives themselves are extended at the higher virus concentrations. It is important to design surface drying experiments for pathogens with realistic levels of input virus, and to consider the role of the carrier and matrix if the results are to be clinically relevant.

**IMPORTANCE:** During the COVID-19 pandemic, a lot of attention has been paid to the environmental decay of SARS-CoV-2 due to proposed transmission of the virus via fomites. However, published experiments have commenced with very high virus titer inoculums, an experimental design not representative of real-life conditions. The study described here evaluated the impact of initial virus titer on environmental decay of an enveloped virus, using a non-pathogenic surrogate for SARS-CoV-2, enveloped bacteriophage Phi6. We establish that higher concentrations of virus can protect the virus from environmental decay, depending on conditions. This has important implications for stability studies of SARS-CoV-2 and other viruses. Our results point to a limitation in the fundamental methodology that has been used to attribute fomite transmission for almost all respiratory viruses.

## INTRODUCTION

Early in the COVID-19 pandemic, there was an intense focus on fomites (i.e., inanimate objects and surfaces), as possible conduits for transmission of the causative agent, SARS-CoV-2. This was because of a widely repeated contention that a person touching a freshly contaminated surface, not washing hands, then quickly touching their mouth, nose or eyes, would lead to self-inoculation of this respiratory virus. Consequently, considerable effort has been made to determine how long the virus remains infectious after being deposited on various surfaces, and what conditions favor or disfavor viability of the virus on these surfaces (Aboubakr et al, 2020; Biryukov et al., 2020; Bonil et al., 2021; Chan et al, 2020; Chin et al, 2020; Kasloff et al., 2021; Kwon et al., 2021; Pastorino et al., 2020; Paton et al., 2021; Riddell et al, 2020; van Doremalen et al, 2020).

In parallel to these studies, workers tested for the presence of viral RNA on surfaces in hospitals treating COVID-19 patients (Chia et al., 2020; Guo et al., 2020; Ong et al., 2020; Piana et al., 2021; Santarpia et al., 2020). These RT-PCR tests found viral RNA to be present on many surfaces (but did not test for infectious virus, with one exception), and reinforced the perception that fomites were indeed a significant risk factor for transmission of the disease.

In July 2020, one of us published (online) a Comment arguing that the risk of transmission of SARS-CoV-2 by fomites was exaggerated (Goldman, 2020). New information that appeared since then has strengthened this conclusion (Goldman, 2021). The basis for the argument was that the amounts of virus used in experiments for determining how long infectious virus remains viable on surfaces were orders of magnitude too large compared to what someone would actually encounter in a real-world situation. Since the virus decays with a defined half-life depending on the surface, the larger the inoculum, the more half-lives have to be gone through before there is less than one infectious virus particle remaining on the surface. Smaller, more realistic inoculums would survive through fewer half-lives, and therefore much less time would pass before the surface would be free of infectious virus.

Among conditions favoring virus survival on surfaces was the observation that Bovine Serum Albumin (BSA) protected the virus from environmental decay, and extended the time that the virus remained viable (Pastorino et al., 2020). Similar observations with Bovine Serum Albumin were also noted in experiments assessing viability of bacteriophage MS2, and enveloped bacteriophage Phi6 in droplets (Lin et al., 2020). Bacteriophage Phi6 has been considered as a potential non-pathogenic surrogate for enveloped viral pathogens like SARS-CoV-2 and Ebola virus (Fedorenko et al., 2020; Whitworth et al., 2020).

The fact that Bovine Serum Albumin protected at least three viruses (SARS-CoV-2, phage MS2 and phage Phi6) from environmental decay made us wonder if higher concentrations of a virus itself might similarly protect the virus from decay. Indeed, Marr and coworkers suggested the value of “investigating the role of viral titer, which might affect aggregation and other characteristics, on virus survival” (Lin et al., 2020).

There was already a suggestion that this might be the case for SARS-CoV-1. Table 1 in Lai et al (2005) indicated that survival of SARS on paper, cotton gowns, and disposable gowns was much greater for a 10^6^ inoculum compared to 10^4^. At a 10^4^ inoculum, infectious virus was not detectable after 5 minutes, but with a 10^6^ inoculum, infectious virus remained detectable for 24 hours. This result suggested that the virus half-life was greatly extended with higher amounts of input virus.

**TABLE 1.** Extent of survival of Phi6 dried on plastic is increased at higher phage concentrations

| <u>Line</u> | <u>Time dry (min)</u> | <u>Inoculum</u> | <u>Recovered</u> | <u>% Recovered</u> | <u>Half-life (min)</u> |
| --- | --- | --- | --- | --- | --- |
| 1 | 0 | $6 \times 10^3$ | $5.6 \times 10^3$ | 93 | |
| 2 | 0 | $6 \times 10^2$ | $4.2 \times 10^2$ | 70 | |
| 3 | 0 | 60 | 1 | 2 |  |
| 4 | 0 | 6 | 0 | 0 |  |
| 5 | 15 | $1.2 \times 10^4$ | $1 \times 10^4$ | 83 | 57 |
| 6 | 15 | $1.2 \times 10^3$ | $3.2 \times 10^2$ | 27 | 8 |
| 7 | 15 | $1.2 \times 10^2$ | 13 | 11 | 5 |
| 8 | 15 | 12 | 0 | 0 |  |
| 9 | 30 | $4.7 \times 10^4$ | $2 \times 10^4$ | 43 | 24 |
| 10 | 30 | $4.7 \times 10^3$ | $7.6 \times 10^2$ | 16 | 11 |
| 11 | 30 | $4.7 \times 10^2$ | 18 | 4 | 6 |
| 12 | 30 | 47 | 0 | 0 |  |
| 13 | 60 | $3.2 \times 10^4$ | $1.2 \times 10^4$ | 38 | 42 |
| 14 | 60 | $3.2 \times 10^3$ | $1.4 \times 10^2$ | 4 | 13 |
| 15 | 60 | $3.2 \times 10^2$ | 0 | 0 | |
| 16 | 60 | 32 | 0 | 0 |  |

Amounts of phage Phi6 as shown in the column "Inoculum" were dried in polypropylene tubes.
Samples were reconstituted in 100 μl saline at the times after drying shown in the table. The
amounts of viable phage remaining in the reconstituted samples were determined and shown in the
column "Recovered". Half-life was calculated as described in Materials and Methods.

In the work reported here, we have investigated the role of initial virus concentration on environmental decay of phage Phi6. We assayed survival of virus samples dried in plastic tubes for various lengths of time after drying. We show that higher input virus concentrations do indeed exhibit significantly larger percent survival and longer half-lives compared to lower virus input, however this effect is influenced by the inoculating matrix. The protective effect of higher virus concentrations also was observed for virus samples kept in solution at 4°C. This protective effect was not found for virus placed in Luria-Bertani (LB) growth medium (which contains Tryptone and Yeast Extract) before being placed in plastic tubes for drying.

## MATERIALS AND METHODS

### Phage preparation

Bacteriophage Phi6, and a bacterial host strain that it grows in, *Pseudomonas syringae* var phaseolicola HB10Y, were generous gifts of Lenny Mindich (now retired) of the Public Health Research Institute of Rutgers University. Cells grown overnight in LB medium in tubes shaken at 25°C were used in plaque assays with serial dilutions of virus to obtain countable numbers of plaques. Plaques assays were performed as described in Goldman (2015) except that the plates were incubated at room temperature (approximately 20°C). Luria-Bertani (LB) medium contained 10 g/l Tryptone, 5 g/l yeast extract, 10 g/l NaCl, and NaOH to adjust pH to 7.0 (https://asm.org/getattachment/5d82aa34-b514-4d85-8af3-aeabe6402874/LB-Luria-Agar-protocol-3031.pdf).

Phage stocks were obtained by harvesting the top agar (6.5 g/l in LB) from one or two petri dishes (containing 10 g/l agar in LB) exhibiting confluent lysis of the bacterial lawn. Saline solution (9 g/l NaCl), 1 ml per plate, was added to the top agar, which was transferred to centrifuge tubes and centrifuged at 20,000 x *g* for 5 minutes to remove agar and debris. This supernatant, stored at 4°C, comprised the initial phage stock.

### Preparation of phage stock in saline solution

An Amicon Ultra 100K filter device from Millipore was pre-rinsed with 4 ml of distilled water, followed by subsequent washes with 70% ethanol and sterile saline solution, using centrifugation at 4°C in a fixed angle rotor at 5000 x *g*. Up to 4 ml of Phi6 initial phage stock was loaded on this filter unit and centrifuged such that 200 µl volume remained above the filter (typical spin time 15-20 min). Fluid below the filter was discarded, the filter unit was refilled with 3.8 ml of sterile saline solution and the same centrifugation steps were repeated 5 times. The sample (saline stock) was recovered in 200-400 µl volume and virus titer was determined by plaque assay.

### Phage survival following drying in polypropylene tubes

An aliquot from the saline stock was subjected to a sequential series of 10-fold dilutions in saline. Five µl of the stock, and 5 µl of each of the 10-fold dilutions, were placed near the bottom of 1.5 ml capacity conical polypropylene Eppendorf microcentrifuge tubes (Corning). In our early experiments, we allowed samples to air dry but switched to desiccation to save time. Samples were desiccated under house vacuum (approximately 20 mm-Hg) and removed from the desiccator when visually dry. Generally, this took between 15-20 minutes, with higher concentrations of phage exhibiting shorter drying times, except for samples dried in LB medium where all samples took 19-20 minutes to dry. We observed that there were no significant differences in patterns of phage survival between air drying and desiccation. The data reported here were obtained from desiccated samples.

Ambient room humidity was not controlled and varied in the building with a range between 10% to 45% over a period of six months, depending on the weather. However, for most of the experiments reported here, humidity was around 15-25%. Humidity was monitored on a Holmes HHG-150 Comfort Check Hygrometer & Thermometer. Although humidity is known to significantly affect environmental decay of Phi6 (Fedorenko et al, 2020; Lin et al, 2020; Whitworth et al, 2020), all samples within a given experiment were subject to the same humidity, and our interest was only to ascertain effects of initial viral concentration. Also, experiments measuring virus survival in real world conditions, as has been done for SARS-CoV-2 (e.g., Mondelli et al, 2020; Ben-Shmuel et al, 2020), do not control for humidity, which is variable. Ambient room temperature was also not controlled, but generally was maintained around 20°C.

Dried samples were reconstituted with 100 µl saline added to the Eppendorf tubes and vortexed. The titer of viable virus remaining in each tube was then determined by plaque assay. Half-life for the virus in a particular sample was calculated using the tool at: https://www.calculator.net/half-life-calculator.html?type=1&nt=25&n0=2300&t=60&t12=&x=45&y=11

All experiments were repeated 1 – 3 times; representative experiments are shown in the tables. Because of variations in conditions from experiment to experiment (such as humidity and initial titer of phage stock), we did not pool results to obtain averages. However, the relative percent survival in each experiment was generally consistent for the same time points, and the patterns were reproducible within experimental limits.

For experiments with LB medium, the initial phage stock was subjected to a sequential series of 10-fold dilutions in LB medium. The remainder of the protocol was the same as above.

### Phage survival in solution

The serial dilutions used for the dry time experiments were stored at 4°C in the dark for later testing. After the number of days as shown in the tables, the titer of the phage in each dilution was determined by plaque assay, and compared to the initial titers as measured on day 1.

## RESULTS AND DISCUSSION

### Survival of Phi6 dried on plastic is increased at higher phage concentrations

Table 1 shows survival of Phi6 dried in plastic tubes and left for various lengths of times as a function of initial virus concentration. The simple act of drying the phage led to loss of nearly all viable phage at the lower phage input concentrations (lines 3 and 4), while having no significant effect (93% recovery) on the highest phage concentration tested (line 1), and a small effect (70% recovery) on a 10-fold lower initial phage concentration (line 2).

Similar patterns of protection by higher initial phage concentrations were also seen for all subsequent lengths of time the phage remained dry in the tube. At 15 minutes dry time, we began to see some loss of survival from the most concentrated initial virus input (line 5, 83% recovery) compared to a 10-fold lower virus input (line 6, 27% recovery), As was seen for the samples dried and assayed immediately, the lowest virus inputs led to loss of almost all viable phage (line7, 11% recovery, line 8, none recovered).

After 30 or 60 minutes of dry-time, the highest initial phage inputs began to show significant environmental decay, with just 43% recovery after 30 minutes (line 9), and 38% recovery at 60 minutes (line 13). But even more substantial environmental decay was observed for the lower input virus samples (Lines 10-12 for 30-minute dry time, lines 14-16 for 60 minutes).

For those samples with measurable virus survival, we were able to calculate the half-lives of virus in those samples (Table 1), which were commensurate to the percent survival observed. That is, at the higher phage initial input levels, half-lives were much longer than the half-lives at lower initial phage input, e.g., compare line 5 (57 minute half-life) to line 6 (8 minute half-life), or line 9 (24 minute half-life) to line 11 (6 minute half-life), or line 13 (42 minute half-life) to line 14 (13 minute half-life).

The results in Table 1 show that the higher phage input concentrations delay and protect dried phage from environmental decay compared to lower phage input concentrations. This effect is also visualized in Figure 1A. The black circles show the relationship between the log PFU (plaque forming units) inoculated onto the surface versus the level recovered. The dashed line represents the best fit regression line. The solid line represents a relationship where all inoculated viruses would be recovered (i.e., 100% recovery). It’s clear from the difference between the slopes of the two lines that recovery is progressively lower as the inoculation level declines. In those experiments where inoculated virus was not recovered, no recovery (0 PFU) is visualized in the figure at −1 log PFU.

**Figure 1.**
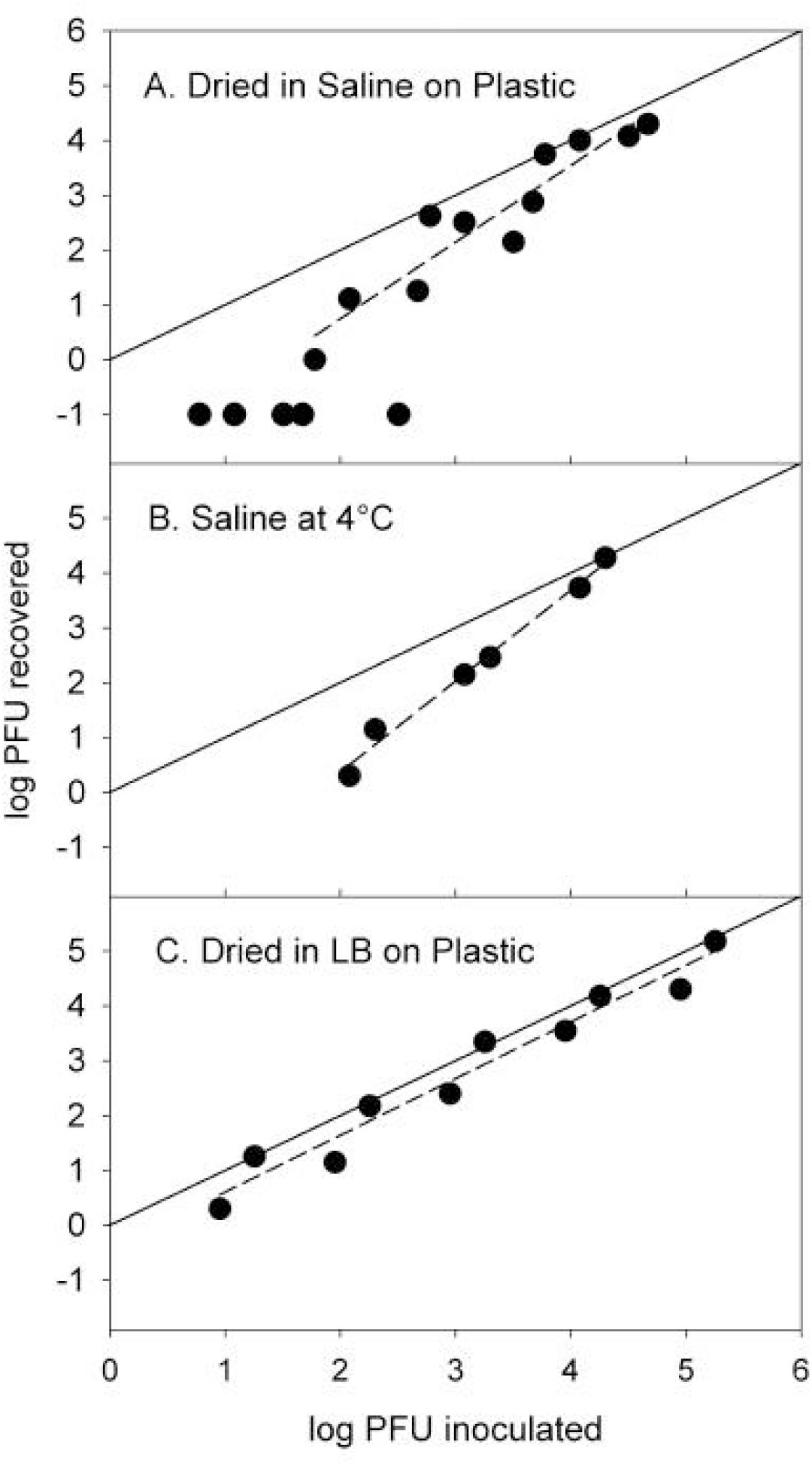
Relationship between the log PFU inoculated versus the level recovered (black circles) for (A) bacteriophage phi 6 on plastic in saline at room temperature, (B) bacteriophage phi 6 in saline at 4°C and (C) bacteriophage phi 6 on plastic in Luria-Bertani broth at room temperature. The dashed lines represent the best fit regression line for each dataset. The solid line represents the line of perfect recovery where all inoculated viruses would be recovered (i.e., 100% recovery). Samples in those experiments where inoculated virus was not recovered (0 PFU) are visualized in the figure at −1 log PFU.

### Survival of Phi6 in saline at 4°C is increased at higher phage concentrations

We decided to test whether there was an effect of phage concentration on virus stability in solution, as was seen for phage dried on surfaces. Table 2 shows little decay of phage in solution at 4°C at the highest concentrations tested after 20 days (line 1), or about half decay after 56 days (line 4). But as the input concentration is reduced, decay increases at both time points (lines 2 and 5), with percent recovery in single digits for the lowest input phage samples (lines 3 and 6). These results can also be visualized in Figure 1B. As in Figure 1A, the black circles show the relationship between the log PFU inoculated onto the surface versus the level recovered. The dashed line represents the best fit regression line, and the solid line represents a relationship where all inoculated viruses would be recovered (i.e., 100% recovery). As in Figure 1A, it’s clear from the difference between the slopes of the two lines that recovery is progressively lower as the inoculation level declines. These experiments had no trials where inoculated virus was not recovered.

**TABLE 2.** Survival of Phi6 in saline at 4°C is increased at higher phage concentrations

| <u>Line</u> | <u>Time (days)</u> | <u>Initial input</u> | <u>Recovered</u> | <u>% Recovered</u> | <u>Half-life (days)</u> |
| --- | --- | --- | --- | --- | --- |
| 1 | 20 | $2 \times 10^4$ | $1.9 \times 10^4$ | 95 | 270 |
| 2 | 20 | $2 \times 10^3$ | $2.9 \times 10^2$ | 15 | 7 |
| 3 | 20 | $2 \times 10^2$ | 14 | 7 | 5 |
| 4 | 56 | $1.2 \times 10^4$ | $5.4 \times 10^3$ | 45 | 49 |
| 5 | 56 | $1.2 \times 10^3$ | $1.4 \times 10^2$ | 12 | 18 |
| 6 | 56 | $1.2 \times 10^2$ | 2 | 2 | 9 |

Amounts of phage Phi6 in saline shown in the column "Initial input" were placed in tubes on day 1
and kept at 4°C for the times indicated, at which point the amounts of viable phage remaining in
the tubes were determined and shown in the column "Recovered". Half-life was calculated as
described in Materials and Methods.

### Protection of Phi6 from decay by LB medium

In our earliest experiments, we used the “initial phage stock” (see Materials and Methods), which did show protection from decay at the higher phage concentrations. But we realized that this stock also had some level of LB medium present, derived from the soft top agar layer containing the lysate in phage preparation. Therefore, dilutions of the initial phage stock also diluted part of the medium the phage had been grown in, which is what prompted us to develop the filtration method described in the Materials and Methods. This allowed the phage to be tested without residual components of the growth medium, which could affect phage survival.

Nevertheless, we wondered whether growth medium itself might also protect the phage from decay, similar to observations with BSA (Lin et al., 2020). Table 3 shows that this is indeed the case. When phage samples were diluted in LB medium instead of saline, phage were essentially completely protected from decay following a half hour dry time (lines 1-5). This is in marked contrast to the results when phage were diluted in saline (Table 1). After extending the dry time for LB-containing samples to 24 hours, environmental decay was now evident in the samples, with recoveries of phage ranging between 16-39% (lines 6-10). There was no effect of varying the amount of initial input phage, showing that LB medium delays and protects even lower concentrations of phage from environmental decay. This is evident from Figure 1C, which shows that as the level of log PFU inoculated declines, the log PFU declines proportionally, and the slope of the regression line (dashed line) is essentially parallel to the line of 100% recovery (solid black line). The regression lies slightly under the line of 100% recovery, which indicates that most, but not all the virus inoculated was recovered with the same rate of environmental decay, but this varied slightly from experiment to experiment. These experiments also had no trials where inoculated virus was not recovered.

**TABLE 3.**
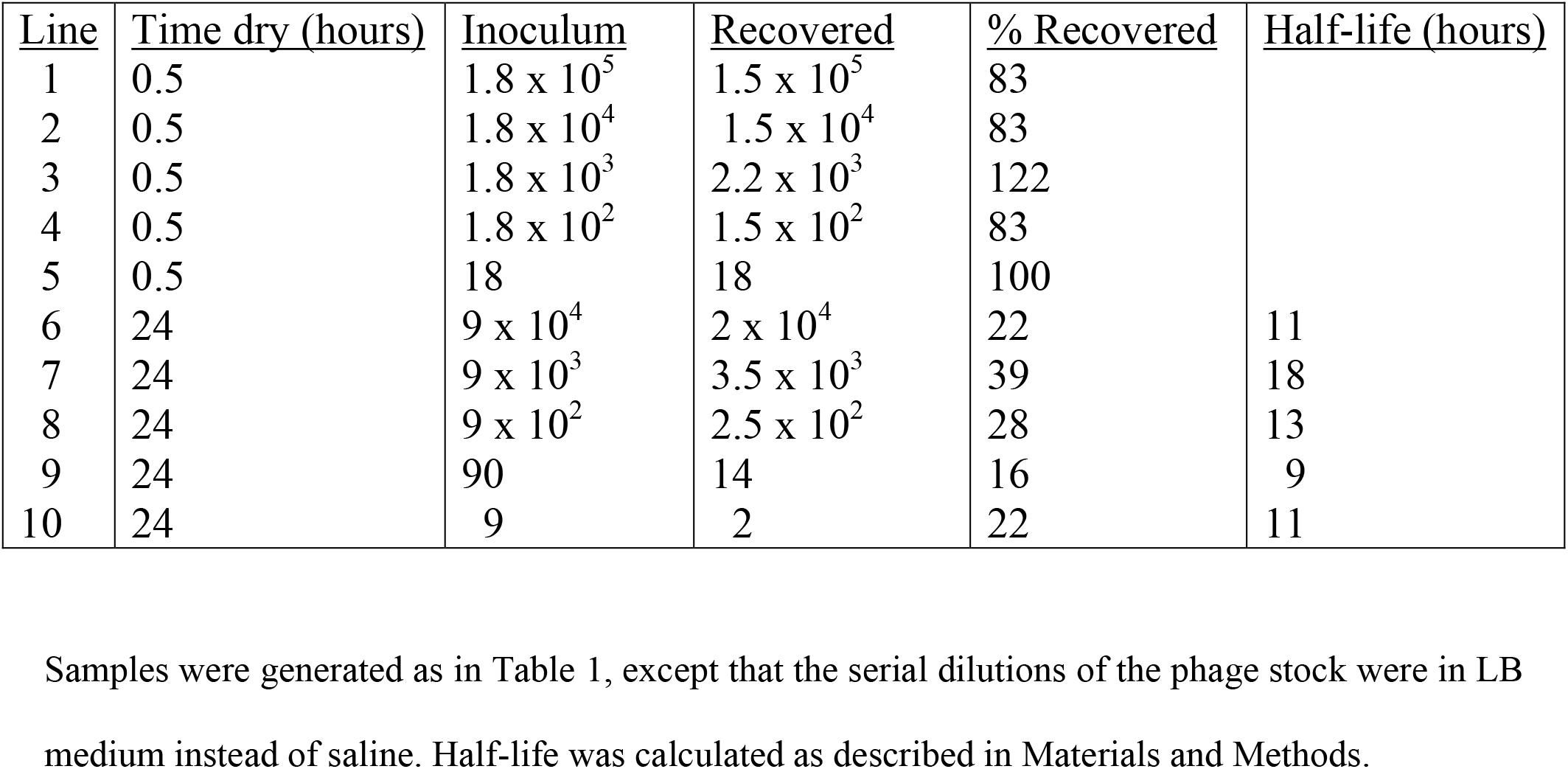
Protection of Phi6 from decay by LB medium dried on plastic

| <u>Line</u> | <u>Time dry (hours)</u> | <u>Inoculum</u> | <u>Recovered</u> | <u>% Recovered</u> | <u>Half-life (hours)</u> |
| --- | --- | --- | --- | --- | --- |
| 1 | 0.5 | $1.8 \times 10^5$ | $1.5 \times 10^5$ | 83 | |
| 2 | 0.5 | $1.8 \times 10^4$ | $1.5 \times 10^4$ | 83 | |
| 3 | 0.5 | $1.8 \times 10^3$ | $2.2 \times 10^3$ | 122 | |
| 4 | 0.5 | $1.8 \times 10^2$ | $1.5 \times 10^2$ | 83 | |
| 5 | 0.5 | 18 | 18 | 100 |  |
| 6 | 24 | $9 \times 10^4$ | $2 \times 10^4$ | 22 | 11 |
| 7 | 24 | $9 \times 10^3$ | $3.5 \times 10^3$ | 39 | 18 |
| 8 | 24 | $9 \times 10^2$ | $2.5 \times 10^2$ | 28 | 13 |
| 9 | 24 | 90 | 14 | 16 | 9 |
| 10 | 24 | 9 | 2 | 22 | 11 |

Samples were generated as in Table 1, except that the serial dilutions of the phage stock were in LB
medium instead of saline. Half-life was calculated as described in Materials and Methods.

Our data demonstrate that bacteriophage Phi6, an enveloped virus that has been considered a potential non-pathogenic surrogate for SARS-CoV-2 transmission studies, exhibits a slower loss of infectivity from environmental decay by higher initial virus concentrations, depending upon the carrier. This slower loss of infectivity is true both when the phage are dried on plastic surfaces, and when the phage are left in saline solution in the refrigerator. The protection at higher phage concentrations is reflected in longer half-lives compared to lower phage concentrations.

Phage survival on dried surfaces can be greatly affected by the type of surface, by temperature and humidity, and by the medium containing the phage (Lin et al., 2020; Fedorenko et al., 2020; Whitworth et al., 2020). Indeed, we observed dramatic delay and protection of phage by LB growth medium, which superseded the effects of initial phage concentration. Thus, an important limitation of our results is that we do not know what effect (if any) on phage survival would result from other natural additions, such as mucous for example, which has been tested for SARS-CoV-2 (Matson et al., 2020).

A recent study compared environmental stability of dried SARS-CoV-2 for two different initial virus inocula (4 × 10^5^ versus 4 × 10^3^) on stainless steel, and did not observe additional protection in the rate of decay at the higher concentration (Paton et al., 2021). The SARS-CoV-2 samples used in this study were in growth medium including fetal bovine serum, which may be more comparable to our results for Phi6 in LB medium. Also, their lower tested concentration (4 × 10^3^) may still have been too high to observe accelerated decay at lower virus concentrations.

Our results have implications for experiments measuring SARS CoV-2 survival on surfaces as purported sources of transmission. High initial amounts of input virus on a surface may generate artificially inflated survival times compared to realistic lower levels of virus that a subject would normally encounter, not only because there are extra half-lives to go through at the higher concentrations, but also because the half-lives themselves are extended at the higher virus concentrations.

The implications of our findings are also relevant for experiments supporting transmission of respiratory viruses by fomites in general. With the exception of Respiratory Syncytial virus, the belief that most, if not all, of these viruses can be spread by fomites is based solely on dried virus stability experiments (e.g. Kutter et al, 2018). In the case of rhinovirus, the major cause of the common cold, an early report demonstrating experimental fomite transmission used unrealistic conditions (Gwaltney and Hendley, 1982). A subsequent study closer to real-life conditions disproved this route of transmission, at least to a first approximation (Dick et al, 1987). Apparent low efficiency of virus transfer by fingers also needs to be considered when assessing the possibility of fomite transmission, as contact with hands might inactivate some viruses (Weber and Stilianakis, 2021).

We do not know why higher concentrations of input Phi6 protect the virus from decay, or why it is influenced by the carrier. It could just be a result of higher protein levels in the phage solution buffering the phage particles from the effects of drying out. Marr and coworkers pointed out that as liquid evaporates, the concentration of solutes (in our case, salt) increases in the microenvironment (Lin et al., 2020), which may affect survival. Further, virus aggregation at higher concentrations is likely, and such aggregates may be protective (Gerba and Betancourt, 2017), or simply manifest as protective because clusters of viruses will be counted as single PFU, but behave kinetically as more resistant viruses. Whatever the cause for this protection, it is even more imperative to design surface drying experiments for pathogens with realistic levels of input virus, and account for the effects of carrier and matrix if the results are to be clinically relevant.

## ACKNOWLEDGEMENTS

Funding sources for EG’s laboratory were grants from the New Jersey Alliance for Clinical and Translational Science (NJ ACTS), and the Rutgers Research Council. We are grateful to Matthew Igo for advice and sharing a lab protocol for Phi6 plaque assays. We acknowledge with gratitude the many helpful comments and insights offered by Wlodek Mandecki.

## AUTHOR CONTRIBUTIONS

RB performed the dry time experiments; EG performed the solution experiments with assistance from RB. MC assisted in preparation of materials used and developed the filtration method for generating saline solutions of phage from LB stocks. First draft of the manuscript was prepared by EG, with input from a report by RB. DS generated the figure, and assisted in interpretation of the experiments and revisions of the manuscript. All authors reviewed the manuscript, made corrections, and approved the final version. Overall supervision of the project was under the direction of EG.

## REFERENCES

Aboubakr, HA, Sharafeldin, TA, Goyal, SM. Stability of SARS-CoV-2 and other coronaviruses in the environment and on common touch surfaces and the influence of climatic conditions: A review. Transbound Emerg Dis. 2020; 00: 1– 17. https://doi.org/10.1111/tbed.13707

Ben-Shmuel A, Brosh-Nissimov T, Glinert I, et al. Detection and infectivity potential of severe acute respiratory syndrome coronavirus 2 (SARS-CoV-2) environmental contamination in isolation units and quarantine facilities. Clin Microbiol Infect. 2020; 26:1658–1662. https://doi.org/10.1016/j.cmi.2020.09.004

Biryukov, J., Boydston, J. A., Dunning, R. A., Yeager, J. J., Wood, S., Reese, A. L., Ferris, A., Miller, D., Weaver, W., Zeitouni, N. E., Phillips, A., Freeburger, D., Hooper, I., Ratnesar-Shumate, S., Yolitz, J., Krause, M., Williams, G., Dawson, D. G., Herzog, A., Dabisch, P., … Altamura, L. A. (2020). Increasing Temperature and Relative Humidity Accelerates Inactivation of SARS-CoV-2 on Surfaces. mSphere, 5(4), e00441–20. https://doi.org/10.1128/mSphere.00441-20

Bonil, L.; Lingas, G.; Coupeau, D.; Lucet, J.-C.; Guedj, J.; Visseaux, B.; Muylkens, B. Survival of SARS-CoV-2 on Non-Porous Materials in an Experimental Setting Representative of Fomites. Coatings 2021, 11, 371. https://doi.org/10.3390/coatings11040371

Chan KH, Sridhar S, Zhang RR, et al. (2020) Factors affecting stability and infectivity of SARS-CoV-2. J Hosp Infect;106(2):226–231. https://doi.org/10.1016/j.jhin.2020.07.009

Chia PY, Coleman KK, Tan YK, et al. Detection of air and surface contamination by SARS-CoV-2 in hospital rooms of infected patients. Nat Commun. 2020;11(1):2800. Published 2020 May 29. https://doi.org/10.1038/s41467-020-16670-2

Chin AWH, Chu JTS, Perera MRA, et al. Stability of SARS-CoV-2 in different environmental conditions. Lancet Microbe. 2020;1(1):e10. https://doi.org/10.1016/S2666-5247(20)30003-3

Dick EC, Jennings LC, Mink KA, Wartgow CD, Inhorn SL. Aerosol transmission of rhinovirus colds. J Infect Dis. 1987; 156: 442–8. https://doi.org/10.1093/infdis/156.3.442

Fedorenko, A., Grinberg, M., Orevi, T. et al.. Survival of the enveloped bacteriophage Phi6 (a surrogate for SARS-CoV-2) in evaporated saliva microdroplets deposited on glass surfaces. Sci Rep 10, 22419 (2020). https://doi.org/10.1038/s41598-020-79625-z

Gerba CP, Betancourt WQ. Viral Aggregation: Impact on Virus Behavior in the Environment. Environ Sci Technol. 2017 Jul 5;51(13):7318–7325. https://doi.org/10.1021/acs.est.6b05835

Goldman E (2015) Plaque Assay for Bacteriophage. In Practical Handbook of Microbiology, 3rd Edition (Goldman E and Green L, eds.), CRC Press, Boca Raton, FL, pp. 93–97.

Goldman E. Exaggerated risk of transmission of COVID-19 by fomites. Lancet Infect Dis. 2020; 20:892–893. https://doi.org/10.1016/S1473-3099(20)30561-2

Goldman E. SARS Wars: the fomites strike back. Appl Environ Microbiol. 2021 Apr 30:AEM.00653-21. https://doi.org/10.1128/AEM.00653-21.

Guo Z, Wang Z, Zhang S, et al. Aerosol and Surface Distribution of Severe Acute Respiratory Syndrome Coronavirus 2 in Hospital Wards, Wuhan, China, 2020. Emerging Infectious Diseases. 2020;26(7):1583–1591. https://doi.org/10.3201/eid2607.200885.

Gwaltney JM Jr, Hendley JO. Transmission of experimental rhinovirus infection by contaminated surfaces. Am J Epidemiol. 1982 Nov;116(5):828–33. https://doi.org/10.1093/oxfordjournals.aje.a113473

Kasloff SB, Leung A, Strong JE, Funk D, Cutts T. Stability of SARS-CoV-2 on critical personal protective equipment. Sci Rep. 2021 Jan 13;11(1):984. https://doi.org/10.1038/s41598-020-80098-3.

Kutter JS, Spronken MI, Fraaij PL, Fouchier RA, Herfst S. Transmission routes of respiratory viruses among humans. Curr Opin Virol. 2018; 28:142–151. https://www.sciencedirect.com/science/article/pii/S1879625717301773?via%3Dihub

Kwon T, Gaudreault NN, Richt JA. Environmental Stability of SARS-CoV-2 on Different Types of Surfaces under Indoor and Seasonal Climate Conditions. Pathogens. 2021 Feb 18;10(2):227. https://doi.org/10.3390/pathogens10020227

Lai MY, Cheng PK, Lim WW. Survival of severe acute respiratory syndrome coronavirus. Clin Infect Dis. 2005; 41: e67–71. https://doi.org/10.1086/433186

Lin K, Schulte CR, Marr LC (2020) Survival of MS2 and Φ6 viruses in droplets as a function of relative humidity, pH, and salt, protein, and surfactant concentrations. PLoS ONE 15(12): e0243505. https://doi.org/10.1371/journal.pone.0243505

Matson MJ, Yinda CK, Seifert SN et al. Effect of Environmental Conditions on SARS-182 CoV-2 Stability in Human Nasal Mucus and Sputum. Emerg Infect Dis. 2020; 26: 2276-8. 183 https://dx.doi.org/10.3201/eid2609.202267

Mondelli MU, Colaneri M, Seminari EM, et al. Low risk of SARS-CoV-2 transmission by fomites in real-life conditions Lancet Infect Dis. 2020; (published online September 29.) https://doi.org/10.1016/S1473-3099(20)30678-2

Ong SWX, Tan YK, Chia PY, Lee TH, Ng OT, Wong MSY, Marimuthu K. Air, Surface Environmental, and Personal Protective Equipment Contamination by Severe Acute Respiratory Syndrome Coronavirus 2 (SARS-CoV-2) From a Symptomatic Patient. JAMA. 2020 Apr 28;323(16):1610–1612. https://doi.org/10.1001/jama.2020.3227.

Pastorino B, Touret F, Gilles M et al. Prolonged Infectivity of SARS-CoV-2 in Fomites. Emerg Infect Dis. 2020; 26: 2256–7. https://dx.doi.org/10.3201/eid2609.201788

Paton S, Spencer A, Garratt I, Thompson K-A, Dinesh I, Aranega-Bou P, Stevenson D, Clark S, Dunning J, Bennett A, Pottage T. Persistence of SARS-CoV-2 virus and viral RNA in relation to surface type and contamination concentration. Appl Environ Microbiol. May 2021, AEM.00526-21; https://aem.asm.org/content/aem/early/2021/05/03/AEM.00526-21.full.pdf

Piana, A., Colucci, M. E., Valeriani, F., Marcolongo, A., Sotgiu, G., Pasquarella, C., Margarucci, L. M., Petrucca, A., Gianfranceschi, G., Babudieri, S., Vitali, P., D’Ermo, G., Bizzarro, A., De Maio, F., Vitali, M., Azara, A., Romano, F., Simmaco, M., & Romano Spica, V. (2021). Monitoring COVID-19 Transmission Risks by Quantitative Real-Time PCR Tracing of Droplets in Hospital and Living Environments. mSphere, 6(1), e01070–20. https://doi.org/10.1128/mSphere.01070-20

Riddell S, Goldie S, Hill A, et al. The effect of temperature on persistence of SARS-CoV-2 on common surfaces. Virol J 2020; 17: 145. https://doi.org/10.1186/s12985-020-01418-7

Santarpia, J.L., Rivera, D.N., Herrera, V.L. et al.. Aerosol and surface contamination of SARS-CoV-2 observed in quarantine and isolation care. Sci Rep 10, 12732 (2020). https://doi.org/10.1038/s41598-020-69286-3

van Doremalen N, Bushmaker T, Morris DH, Holbrook MG, Gamble A, Williamson BN, Tamin A, Harcourt JL, Thornburg NJ, Gerber SI, Lloyd-Smith JO, de Wit E, Munster VJ. (2020) Aerosol and Surface Stability of SARS-CoV-2 as Compared with SARS-CoV-1. N Engl J Med. 382:1564–1567. https://www.nejm.org/doi/full/10.1056/nejmc2004973

Weber TP, Stilianakis NI. Fomites, hands, and the transmission of respiratory viruses. J Occup Environ Hyg. 2021; 18:1–3. https://doi.org/10.1080/15459624.2020.1845343

Whitworth C, Mu Y, Houston H, Martinez-Smith M, Noble-Wang J, Coulliette-Salmond A, Rose L. 2020. Persistence of bacteriophage phi 6 on porous and nonporous surfaces and the potential for its use as an Ebola virus or coronavirus surrogate. Appl Environ Microbiol 86:e01482–20. https://doi.org/10.1128/AEM.01482-20.

